# Do monkeys dream about food while sleeping?

**DOI:** 10.64898/2026.09.30.755371

**Authors:** Jason Joby, Sini Simon, Thomas Cherian, Georgin Jacob, Jhilik Das, Surbhi Munda, Shubhankar Saha, Debdyuti Bhadra, SP Arun

**Affiliations:** Centre for Neuroscience, Indian Institute of Science, Bangalore 560012; Indian Institute of Science Education and Research, Pune 411008

**Author notes:** indicate equal contributions (Thomas Cherian, Georgin Jacob, Jhilik Das, Surbhi Munda, Shubhankar Saha).

## Abstract

Can we tell what animals are dreaming about from their brain activity? This fascinating question has been explored extensively using the rodent hippocampus, where place cells are sequentially reactivated during sleep, consistent with replay of paths explored while awake. However, dreams involve vivid visual experiences, and whether analogous replays occur in the visual cortex has not been explored, that too in primates. To address this gap, we performed wireless neural recordings from the high-level visual (IT) and motor regions (PMv, PFC) in two monkeys while they viewed food images on a screen while awake, and afterwards while they slept in their natural setting. Decoders trained on sleep/wake neural activity in each region were able to accurately predict the ongoing sleep/wake state, as well as upcoming transitions between states. Interestingly, decoders trained to categorize neural responses to food (but not faces/objects) showed increased probability during REM-like sleep compared to NREM or awake states. Thus, behaviorally relevant visual stimuli seen while awake are reactivated during REM-like sleep. In other words, these monkeys saw food, faces and objects while awake, and dreamt selectively about food while sleeping.

**SIGNIFICANCE STATEMENT:** Dreams involve vivid visual experiences that are often related to what we experience during the day. Understanding the underlying neural basis in primates has been challenging because most brain recordings are performed under artificial conditions in which animals do not even sleep. Here, we recorded wirelessly from high-level visual and motor regions in freely moving monkeys while they viewed images of food, faces and objects and also while they slept in a natural setting. Neural signatures evoked while viewing food (but not faces/objects) were selectively reactivated during REM-like sleep. Thus, these monkeys probably dreamt about food while sleeping.

## INTRODUCTION

Whether we can decode the contents of dreams from brain activity remains a fascinating question. In humans, where it is possible to obtain subjective reports, brain activity during dreaming can be systematically related to subsequent subjective reports (Horikawa et al., 2013; Schönauer et al., 2017; Siclari et al., 2017). In animals, where subjective reports are impossible, our understanding is based on relating brain activity recorded during sleep with experiences in the awake state. The clearest evidence of reactivation comes from rodents, where place cells active along paths traversed while awake are reactivated during sleep (Wilson and McNaughton, 1994; Louie and Wilson, 2001; Lee and Wilson, 2002). Such replay has since been observed in the rodent visual cortex during navigation (Ji and Wilson, 2007), in motor nuclei of songbirds (Dave and Margoliash, 2000), and even during wakeful resting in rodent visual cortex (Xu et al., 2012) and monkey somatosensory and motor regions (Hoffman and McNaughton, 2002).

There is relatively little investigation of sleep in monkeys, partly because most primate studies involve brain recordings in restraint chairs in which animals rarely sleep. Nonetheless, early studies of monkeys sleeping in a restraint chair characterized the architecture of awake, REM and NREM sleep states as well as the associated local field power (Jouvet, 1965; Bert et al., 1975; González et al., 1979). With the advent of wireless recording technologies, neural responses have been recorded in increasingly natural settings, showing unsupervised clustering of neural states into sleep and wake states across multiple brain regions (Hsieh et al., 2008; Gabbott and Rolls, 2013; Yin et al., 2014; Walker et al., 2021; Bukhtiyarova et al., 2022; Yun et al., 2023). Yet, there have been very few studies relating neural activity during sleep in monkeys with neural activity during awake visual experience. As a result, we do not know whether neurons in high-level visual cortex are reactivated during sleep in a manner systematically related to the visual experiences during the awake state. Addressing this gap will also require recording visual responses during the awake state under controlled conditions, and then testing whether these neurons are reactivated while animals sleep in their natural setting.

### Overview of this study

Here, we investigated this issue through wireless brain recordings in freely moving monkeys while they viewed images during a fixation task, and while they slept in a natural setting. We hypothesized that neural responses to behaviorally relevant items like food (but not other items) would be reactivated during specific stages of sleep. To test this hypothesis, we recorded neural responses while monkeys viewed images of food as well as other objects in a fixation task, and also recorded neural responses during sleep. We recorded from the inferior temporal (IT) cortex, a high-level visual region critical for object recognition and perception (Connor et al., 2007; DiCarlo et al., 2012; Conway, 2018). We also recorded simultaneously from premotor/prefrontal regions (ventral premotor cortex, PMv and ventrolateral prefrontal cortex, PFC), which were implanted to study decision-making in these monkeys while they performed other tasks (Rizzolatti and Luppino, 2001; Schrijver et al., 2024).

**Figure 1.**
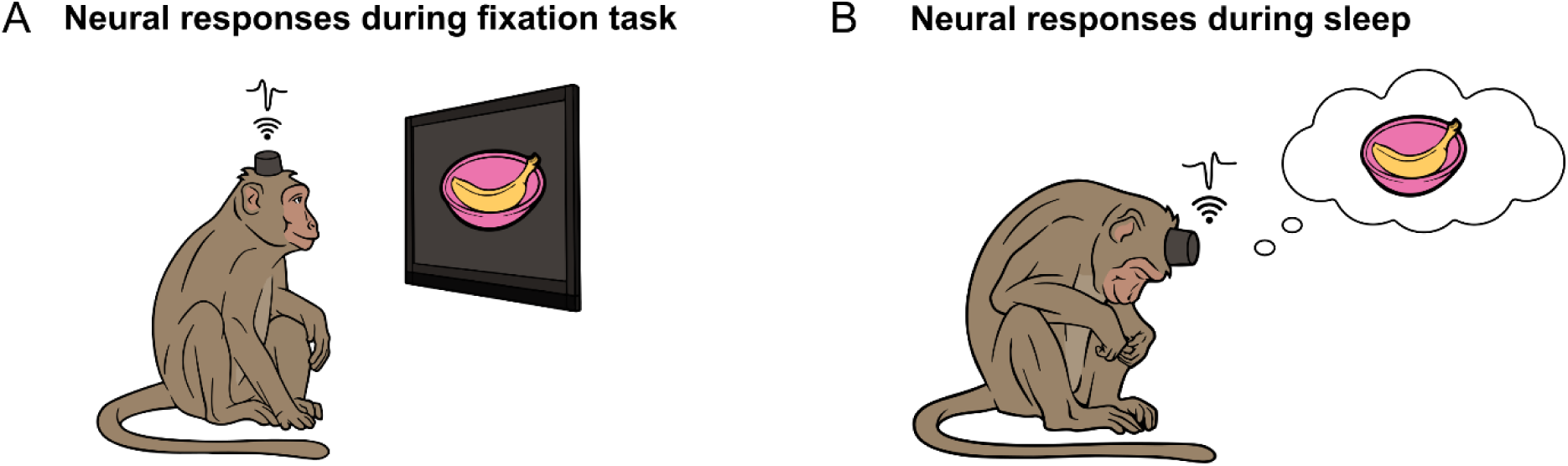
Comparing visual responses during wakefulness and natural sleep. (A) Schematic of a monkey performing a passive fixation task, in which he had to fixate on a series of images to obtain juice reward. These included images of food items that the monkey ate during the awake sessions, as well as images of human experimenters that interacted with him during the day. (B) Schematic of a monkey sleeping in its natural setting. To relate neural responses during sleep with the awake state, we asked whether neural activity decoders trained on food images in the fixation task would reactivate during sleep.

## RESULTS

Each monkey participated in two sessions. In the sleep session, the monkey slept unconstrained in its natural setting at night (Figure 2A). During the day before the sleep session, each monkey performed a fixation task on a touchscreen workstation where it viewed images of food, faces and objects (Figure 2B). During both sessions, we performed wireless brain recordings from 256 electrodes implanted into the higher order visual and motor regions (IT, PMv, PFC) in the two monkeys (Figure 2C; see Methods).

**Figure 2.**
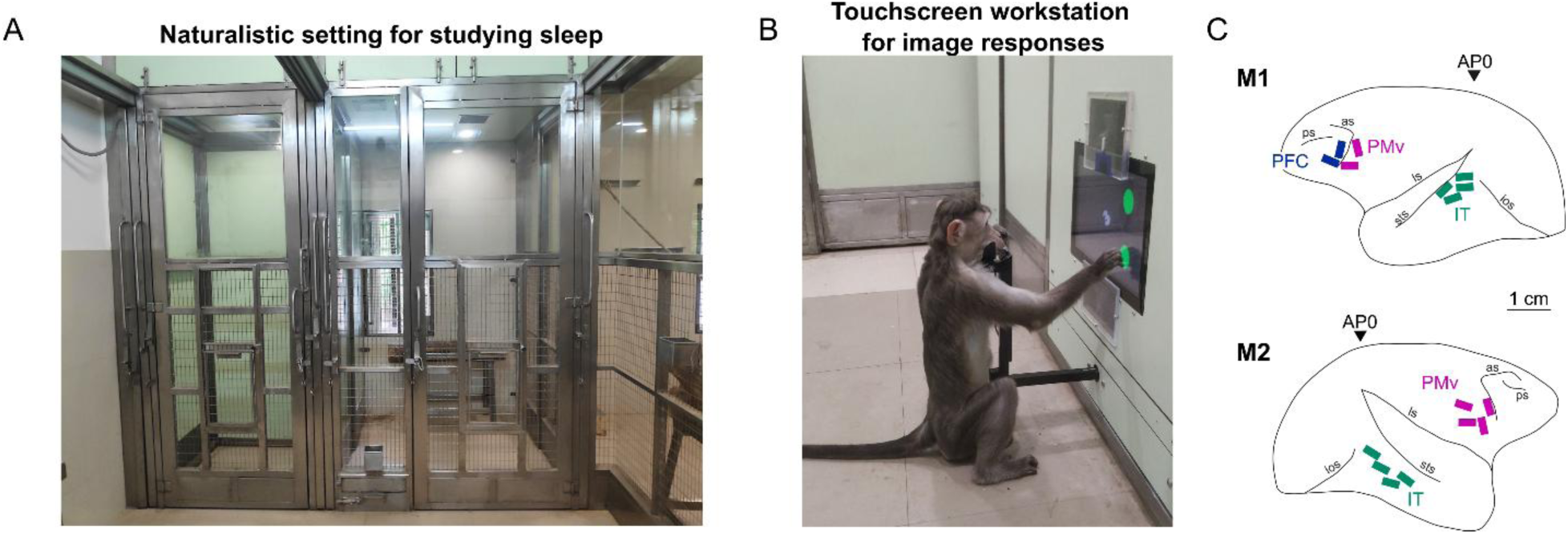
Relating brain activity during sleep & awake states. (A) Naturalistic setting for studying sleep. Each monkey was separated into a large holding partition (*middle*) for monitoring its movements during the sleep session, while its brain activity was recorded wirelessly (Figure panel from Jacob et al, 2021). (B) View of the touchscreen workstation for obtaining neural responses to images, showing a naïve monkey performing a task (Figure panel from Jacob et al, 2021). (C) Brain regions recorded in the two monkeys, reconstructed to scale by aligning sulcal landmarks visible during surgery with MRI scans (Figure panel from Simon et al, 2026).

### Neural signatures of sleep in a natural setting

We began with a detailed characterization of sleep/wake behavior and their neural correlates during the sleep session. We manually annotated the video recordings made during the sleep session in each monkey, with particular emphasis on when the monkey was sleeping or awake. We marked periods when the monkey had his head raised and moving around as “Head Up”. Likewise, periods in which the monkey kept his head down unmoving in a huddled posture, were marked as “Head Down”. Both monkeys spent most of their time sleeping interspersed with brief periods of wakefulness (Figure 3A).

**Figure 3.**
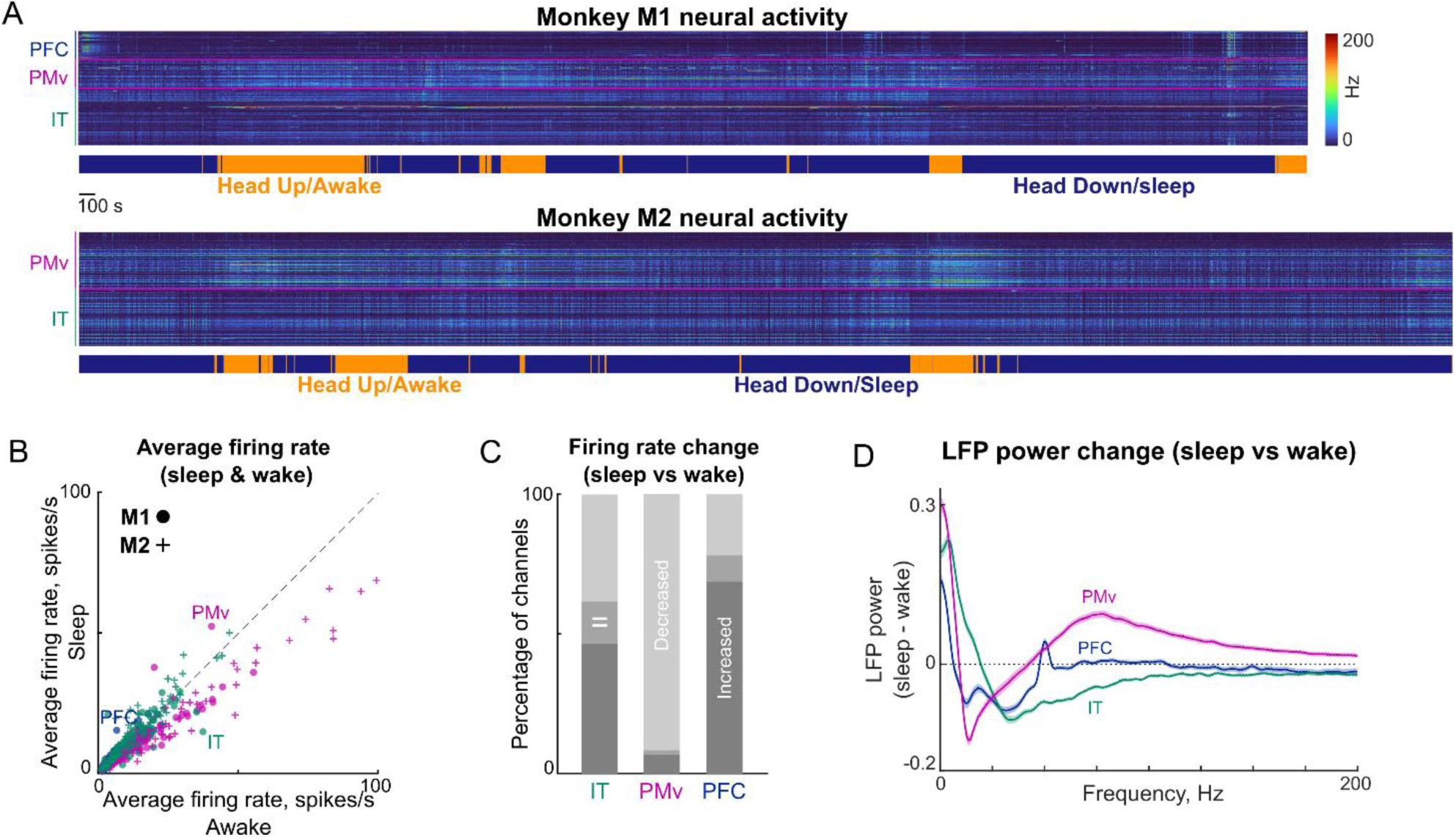
Comparing neural responses during sleep and wake states. (A) Colormap depicting raw firing rates in 1 s bins for all 256 neurons recorded in Monkey M1 during the sleep session. The ethogram below depicts the head up (i.e. awake) and head down (i.e. sleep) states that were manually annotated from the video recording. (B) Average firing rates during wake bins plotted against its average firing rate during sleep bins for neurons in IT (*green*), PMv (*purple*) and PFC (*blue*) from monkey M1 (*circles*) and M2 (*crosses*). (C) Percentage of neurons that showed a significant increase, decrease or no change in firing rate during sleep compared to wake states (p < 0.05, rank-sum test between sleep and wake bins) in IT, PMv and PFC. Most PMv neurons showed lower firing during sleep whereas IT and PFC had roughly equal numbers of neurons with increases and decreases in firing during sleep. (D) Difference in LFP spectral power between sleep and wake states plotted against frequency (mean ± s.e.m across neurons) for IT (*green*), PMv (*purple*) and PFC (*blue*) recording sites. Positive values indicate higher power during sleep. See Figure S1 for a detailed characterization of the LFP signal during sleep and wake states.

Brain activity in both monkeys during the sleep recordings is shown in Figure 3A. It can be seen that brain activity is different in sleep and wake states in both monkeys. To understand the overall response level of each neuron in the sleep and wake states, we plotted the average firing rate across all bins in the sleep and wake states for each neuron (Figure 3B). It can be seen that most PMv neurons have higher firing rates during wake state on average whereas IT & PFC show both increase and decrease in firing rates across neurons (Figure 3C). This is expected, since PMv is a motor region and is likely to be more active when the monkey is moving, such as in the awake state. We also observed characteristic changes in LFP power at the same recording sites: LFP power in the sleep state was higher in low frequencies in all three brain regions (IT, PMv & PFC). Spectral power in PMv was higher in the wake state (∼14 Hz), whereas spectral power in IT was higher in the wake state (∼33 Hz; Figure 3D).

### Decoding of sleep and wake states from neural activity

Next, we asked whether the sleep/wake state could be decoded from neural activity. To this end, we calculated the firing rate in 1 s bins throughout the sleep session and labelled these bins as sleep or wake based on the behavioral state. We then trained a linear decoder on the population firing rate across all recorded neurons across brain regions and asked whether the decoder could correctly identify the behavioral state on unseen time bins. This revealed highly accurate decoding of sleep & wake states (overall accuracy: 93% ± 0.2%; Figure 4A, *left*). Decoders trained on each brain region were also highly accurate (Figure 4A right). Decoding of sleep and wake states was also highly accurate over time during the sleep session (Figure 4B). We obtained similar results using LFP signals (Figure S1).

**Figure 4.**
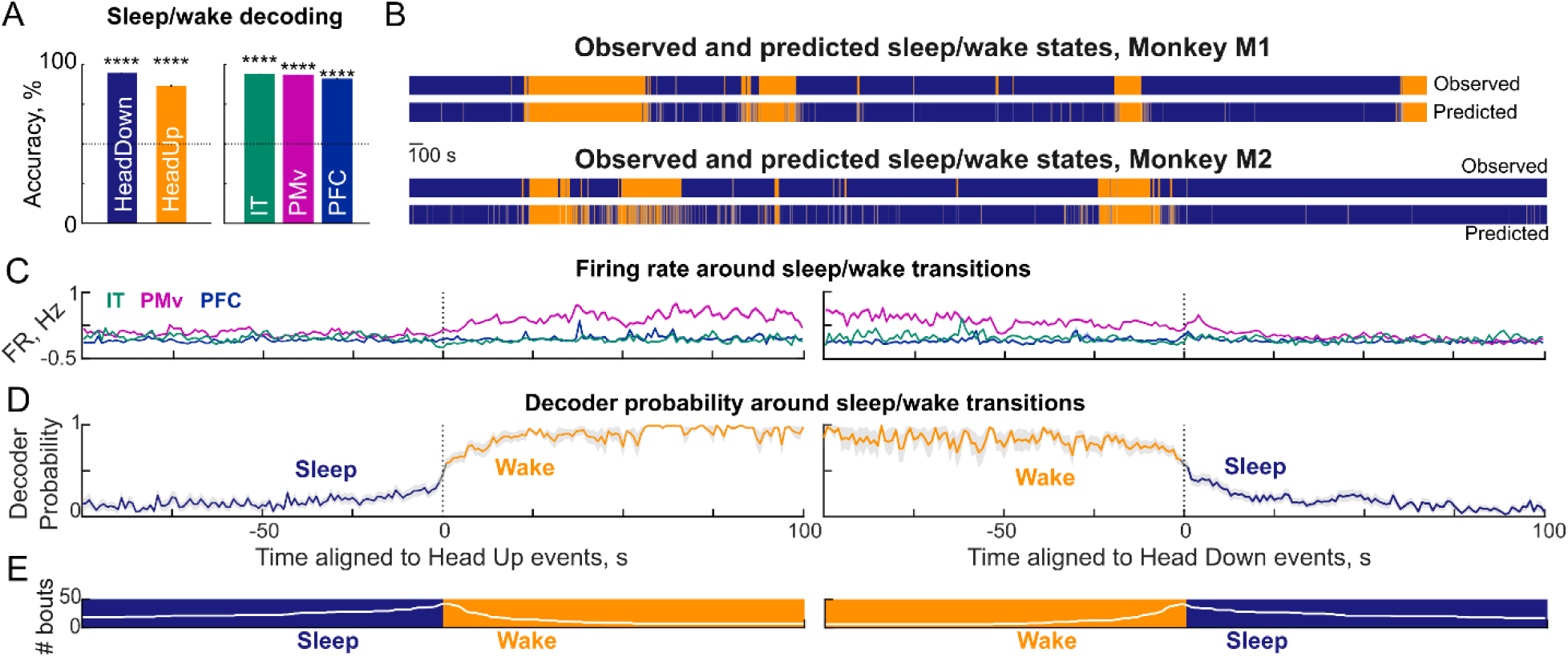
Decoding of sleep and wake states from neural activity. (A) *Left:* Cross-validated decoding accuracy of a linear decoder trained on the population activity in 1 s bins, pooled across monkeys. *Right:* Cross-validated decoding accuracy of decoders trained separately on IT (*green*), PMv (*purple*) and PFC (*blue*). (B) Observed sleep/wake state shown above predicted sleep/wake state from the decoder for each monkey: M1 (*top row*) and M2 (*bottom row*). (C) Firing rates (mean ± s.e.m) in each brain region pooled across two monkeys, relative to the time of sleep/wake state transitions. *Left*: Sleep to wake transitions, showing a clear increase in PMv firing. *Right:* Wake to sleep transitions, showing a clear decrease in PMv firing. (D) Decoder probability (mean ± s.e.m across bouts) for wake state averaged around the time of transitions from sleep to wake (*left*) and wake to sleep (*right*), using all brain regions. (E) Number of bouts contributing at each time to the averaging in panels (D), for sleep to wake (*left*) and wake to sleep (*right*) transitions.

We then wondered what happened around the time of transitions between sleep/wake states. To this end, we calculated the firing rate in each brain region (IT, PMv, PFC) averaged at times relative to each transition from sleep to wake or wake to sleep (Figure 4C). It can be seen that PMv firing rates increased systematically across the transition from sleep to wake and decreased across the wake to sleep transition. By contrast, there was no clear trend in IT & PFC firing rates. Thus, brain activity might reflect impending transitions well before they occur. To confirm this quantitatively, we calculated the average decoder probability for the wake state around times relative to sleep to wake or wake to sleep transitions (Figure 4D). This revealed a gradual rise in the probability of a wake state during the sleep to wake transitions, and conversely a gradual drop during wake to sleep transitions. We obtained similar results using LFP signals (Figure S1).

We conclude that sleep and wake states can be reliably decoded from brain activity in all three brain regions.

### Neural signatures of REM sleep

Having established that brain activity in IT, PMv and PFC changes systematically with the sleep/wake state, we next wondered if any neural signatures during the wake state are reactivated during sleep, particularly during rapid eye movement (REM) sleep. Since we did not have any eye movement or muscle tone data to distinguish REM from non-REM sleep, we adopted an alternative approach to define REM sleep. Since neural activity during REM sleep is similar to the wake state, we calculated the local field potential power (LFP, between 0-50 Hz) in each 1 s time bin for all neurons, and calculated the correlation with the average power during the wake state and with the average power in the sleep state. We then calculated the difference between the correlation between the wake and sleep states for each time bin, and denoted this as LFP similarity (Figure 5A-B). It can be seen that, during certain epochs of the sleep state, neural activity is more similar to the wake state, which we marked as putative REM (pREM). We excluded epochs that were too close to the start of a sleep bout since they might represent transition periods (see Methods). All remaining sleep time bins were marked as NREM (Figure 5A-B). The resulting average LFP similarity for all three bouts showed the expected progression of increasing similarity to the wake state, from NREM to pREM to Awake (NREM, pREM and Awake; Figure 5C).

**Figure 5.**
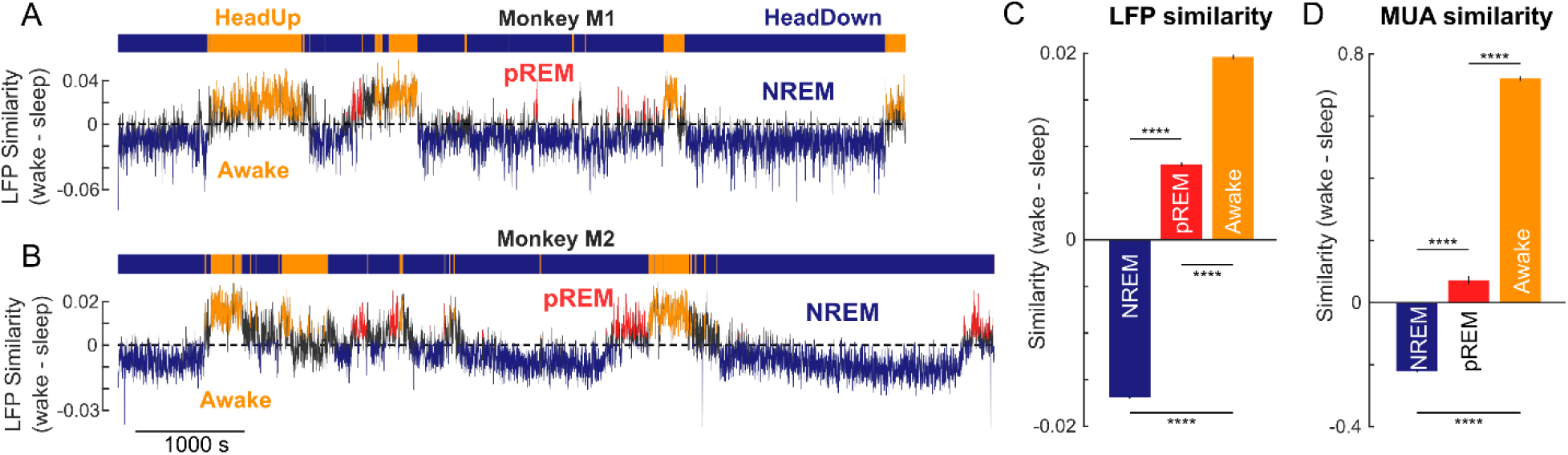
Defining putative REM phases during sleep. (A) LFP similarity (*black*) for monkey M1, calculated as the difference in correlation between the population activity in each 1 s time bin with the average population activity in the wake state minus the sleep state. A positive value of LFP similarity in a time bin indicates that the activity is more similar to the wake state than the sleep state. Consecutive awake bins are marked in *yellow*, putative REM (pREM) bins are marked in *red*, and NREM bins are marked in *blue*. (B) Same as (A) but for monkey M2. (C) Average LFP similarity (wake – sleep) for NREM, pREM and Awake bins pooled across monkeys. Asterisks indicate statistical significance (**** is p < 0.00005, rank-sum test across time bins). (D) Average MUA similarity (wake – sleep) for NREM, pREM and Awake bins defined according to LFP similarity.

To cross-validate our classification of sleep states into NREM and pREM, we calculated a similarity measure based on multiunit activity: as before, we calculated the correlations between the firing rate across all channels with the average firing rate of the wake state and with the average firing rate in the sleep state. The resulting average MUA similarity during the NREM, pREM and awake states are shown for the epochs defined using LFP similarity in Figure 5D. Here too, we see a clear progression of increasing similarity, suggesting that both local field and spiking activity were more similar to the wake state during the pREM epochs.

Thus, brain activity during sleep can be reliably categorized into REM and NREM phases.

### Reactivation of neural signatures of food during REM sleep

Since neural activity in the hippocampus during navigation is reactivated during sleep (Wilson and McNaughton, 1994; Louie and Wilson, 2001; Lee and Wilson, 2002; Ji and Wilson, 2007), we wondered if a similar result could be observed in the visual cortex. During the day of the sleep session, monkeys performed two passive fixation tasks. In the first task, they saw images of food box with and without food pieces (see Methods). In the second task, they saw images of objects and faces (see Methods).

We trained a linear decoder on the trial-wise population activity of IT neurons to images of the box with food and without food and then tested the decoder on the neural activity during the sleep session (Figure 6A). We reasoned that, if monkeys were dreaming about food during the pREM epochs, the decoder probability for food should be higher than NREM or awake epochs. Strikingly, as predicted, the food decoder probability was highest for the pREM epochs compared to NREM, and lowest for the awake epochs (Figure 6B).

**Figure 6.**
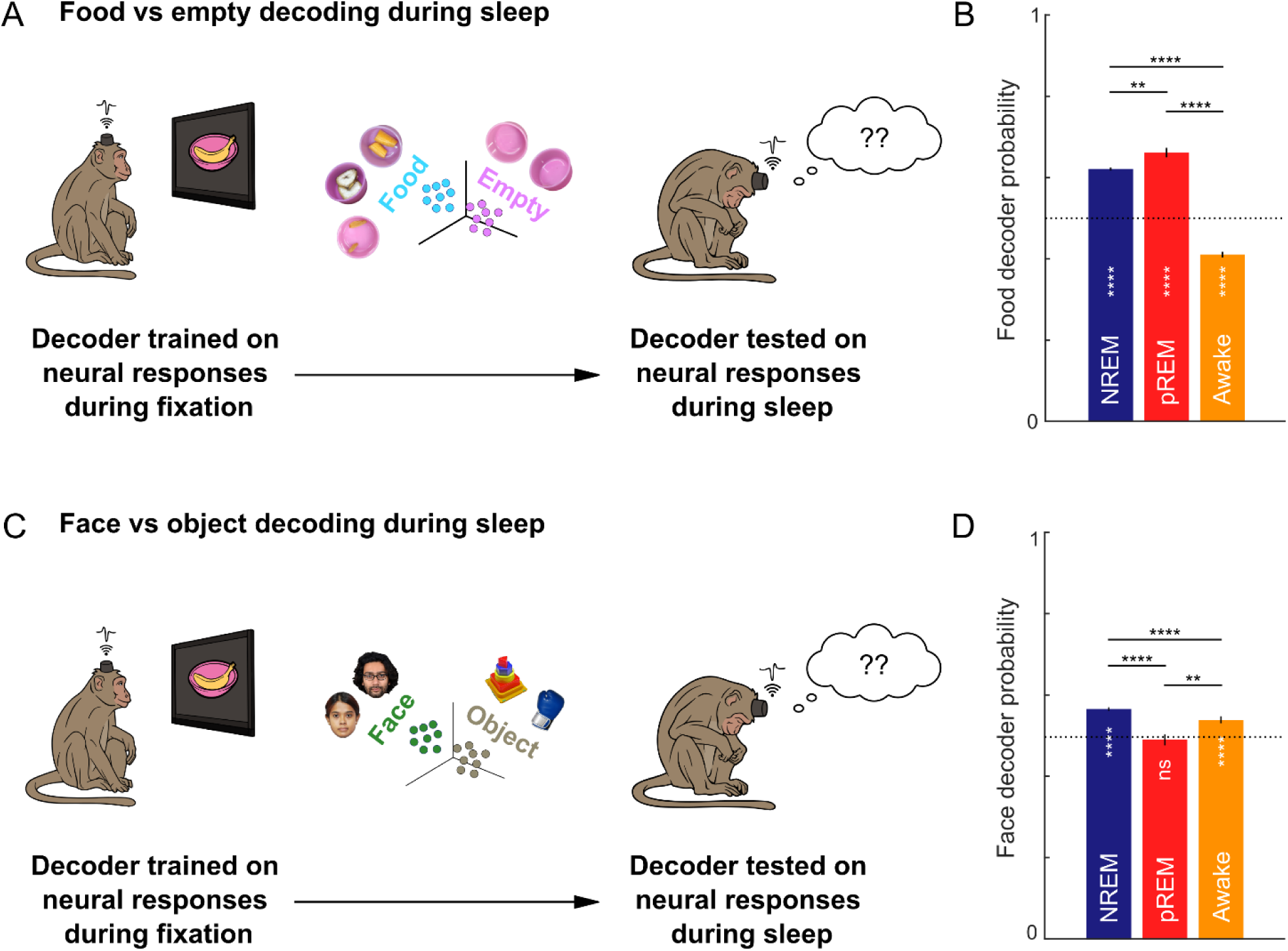
Reactivation of neural signatures of food during sleep. (A) Schematic of food decoding: a linear decoder was trained on neural responses in IT to images of a food box with or without food (*left*) and then tested on neural activity during sleep. (B) Food decoder probability (mean ± s.e.m) in IT for NREM, pREM and Awake time bins pooled across monkeys. It can be seen that food decoding is highest during the pREM bins, followed by NREM bins and least in the awake bins. Asterisks indicate statistical significance comparing brain states (**** is p < 0.0005, rank-sum test comparing decoder probability across time bins). Asterisks within bars indicate statistical significance compared to chance (50%) (**** is p < 0.0005, sign-rank test comparing decoder probability across time bins with that expected by chance). (C) Same as (A) but for decoders trained to distinguish between faces and objects. *[Face images are those of the authors J.D. and S.S.]* (D) Same as (B) but for face decoder probability. It can be seen that face decoding during the pREM bins is no different from chance, but is highest in the NREM phase.

The high food decoding probability during REM sleep could result from neural activity during REM sleep resulting in large, arbitrary projections along the food classifier vector. To rule out this possibility, we projected the neural population vectors during the fixation task as well as during the Awake, NREM and REM epochs in the sleep session along the classifier vector. These magnitudes during sleep were similar to the population vector projections in the fixation task (Figure S2). Thus, food decoding probabilities were not driven by arbitrarily large decoder projections during the sleep session.

If monkeys were really dreaming about food, the food decoder probability should not only be high during REM sleep, but other decoders should have low probability. To investigate this possibility, we trained another decoder on neural responses to objects and faces and tested it on neural responses during sleep (Figure 6C). Indeed, as predicted, the face decoder probability was no different from chance during the pREM phases (Figure 6D). By contrast, it was highest during the NREM bins, followed by the awake bins (Figure 6D). The face decoder also had similar magnitude projections as during fixation (Figure S3). Thus, face decoding probabilities were also not driven by arbitrarily large projections during the sleep session.

We conclude that neural signatures while viewing food (but not face) images in IT neurons are reactivated during REM sleep.

## DISCUSSION

Here, we set out to investigate whether neural signatures elicited in the high-level visual cortex while viewing food are reactivated during sleep. Our main findings are: (1) Neurons show systematic increases and decreases in firing between sleep and wake states; (2) Decoders trained on sleep and wake states can accurately predict not only these states but transitions between these states well in advance; (3) neural signatures elicited by viewing food (but not face) images in the wake state are selectively reactivated during REM-like sleep. These observations suggest that these monkeys dreamt about food (but not faces) while sleeping. Below we discuss our findings in the context of the existing literature.

### Neural signatures of sleep and wake states

Our results reconfirm classic signatures of sleep, such as higher power in low frequencies and lower power in high frequencies (Hsieh et al., 2008; Ishikawa et al., 2017; Bukhtiyarova et al., 2022). However, we have gone further to show that, in all brain regions, there are neurons that are more active during sleep and others that are less active (Figure 3). We have also shown that neural activity can be used to reliably decode the sleep/wake state with high accuracy in all three brain regions (IT, PMv, PFC). This finding is consistent with previous studies showing systematic differences in both neural firing and LFP between sleep and wake states.

Interestingly, we have found that neural decoders trained to distinguish sleep/wake states showed systematic changes around the time of state transitions, often well in advance (Figure 4). The gradual rise and fall of neural decoders suggest that neural activity begins to signal the impending sleep or wake state several seconds in advance, which matches our own subjective experiences of sleeping and waking. The gradual change in decoding probability near sleep/wake transitions could be driven by signals of neural origin driven by sleep centers in the brain, but also by behavioral changes such as the monkey moving around to settle into sleep. Distinguishing between these possibilities would require more fine-grained recording of muscle activity as well as from the sleep centers in the brain, which are interesting possibilities for future work.

### Reactivation of neural signatures during sleep

Our most important finding was regarding the relationship between neural signatures elicited by food images during the wake state with the neural activity in the sleep state. We trained neural decoders to categorize food as present or absent in a box and asked whether these decoders applied to various phases of sleep would show differences in their output. We found that probability of food was highest during the REM phase. We propose that monkeys were dreaming about food during the REM-like states. This is further corroborated by our additional analyses that (1) decoder projections were in the same range during sleep as they were during the fixation task; (2) this was specific to food decoding, since we did not observe the same effect in a face/object decoder.

However, we urge caution with this interpretation because the result is based on decoders that are forced to distinguish only between very few possibilities. A more compelling analysis would be to reconstruct images from neural activity (Dado et al., 2024) and apply this to brain activity during sleep, although this is beyond the scope of the present study. These are interesting possibilities for future work.

## METHODS

All animal experiments were performed according to a protocol approved by the Institutional Animal Ethics Committee of the Indian Institute of Science, Bangalore (CAF/Ethics/750/2020) and by the Committee for the Purpose of Control and Supervision of Experiments on Animals, Government of India (V-11011(3)/15/2020-CPCSEA-DADF). Many experimental procedures are common to previously reported studies from our laboratory (Ratan Murty and Arun, 2015; Jacob et al., 2021), and summarized briefly below.

### Neurophysiology

We implanted two adult monkeys (Ma*caca radiata,* M1: *Co* & M2: *Di*, both aged 9.5 years, 8 arrays in each monkey) using standard neurophysiological procedures reported previously (Ratan Murty and Arun, 2015; Simon et al., 2026).

### Touchscreen workstation

The screen-based tasks were performed by the monkeys in an electromagnetically shielded recording room with a touchscreen station and eye-tracking capability as reported previously (Jacob et al., 2021). To capture the monkey’s movements, we used three cameras (at three viewpoints) aligned with the neural data acquisition system. The specific details of the recording setup are detailed in a recent study (Simon et al., 2026)

### Alignment of sleep video recordings to neural data

The video cameras at the touchscreen workstation were fully aligned with the neural data acquisition system. But the video camera system in the natural area, where the monkey slept during the recording, were not aligned. To achieve this, we resorted to a manual method, as we identified that the touchscreen station had both types of cameras with the full view of the touchscreen - those which are synchronized and those which are not synchronized. Hence, we created a synchronization sequence in which full-screen white images were flashed on the screen. We then used the rise and fall of these flashes on the screen captured by the set of synchronized and non-synchronized cameras to do the alignment.

### Touchscreen tasks

Before the sleep recordings, both monkeys performed a visual task that involved pictures of food items.

#### Eye calibration block

Each task started with a block for calibrating eye position, as detailed previously (Simon et al., 2026). Briefly, the monkey had to fixate successively at four dots appearing on the screen and received a reward for looking at each dot within a certain window of it appearing. The eye data from these trials was used to learn a linear transformation between the raw (x, y) eye signal voltage and the (x, y) visual angles of each dot on the screen.

#### Visual task

Each monkey performed a fixation task to characterize visual responses as described previously (Simon et al., 2026). Briefly, monkeys had to hold a button that appeared on the screen to initiate the trial, and then fixate on a series of 8 images that appeared for 200 ms each with a 200 ms inter-stimulus interval. Each monkey performed two fixation tasks: First they performed a “visual channel mapping” task which involved viewing 16 images (8 faces & 8 objects). This task was used to determine if the multiunit activity recorded on each electrode was visually responsive. Images consisted of 8 isolated faces of human experimenters familiar to the monkeys, and 8 isolated objects all presented after scaling to a longer dimension of 9.2°. Images were randomly interleaved with the constraint that no two successive images in a trial were identical, and in all we collected 16 repetitions of each image. Second, monkeys performed a “foodbox” task that involved viewing images of a plastic box containing three types of food (raisin, carrot, banana) in a pink-colored box. Each food type was shown in 5 views with an isolated black background and also in 3 views with the food pieces scattered on the floor. Images were presented on the screen scaled to a longer dimension of 7.8°, which was roughly the size we estimated the animals would view the food box during real world interactions. Images were randomly interleaved with the same condition as before, and we collected 16 repetitions per image.

Although we used a fairly liberal fixation window (10° for M1, 8° for M2) because we were unsure about fixation quality, our post-hoc analyses of the data revealed that monkeys maintained accurate fixation throughout the trial (standard deviation of eye position during each trial: 0.63° for horizontal and 0.52° for vertical for M1, and 0.45° for horizontal and 0.45° for vertical for M2, averaged across trials; mean jitter in eye position across trials: 0.15° for horizontal and 0.14° for vertical for M1, and 0.13° for horizontal and 0.14° for vertical for M2).

All data analysis in the fixation task were performed on a window 0 to 0.3 s after image onset for food images and 0-0.3 s after image onset for experimenter faces. Varying these choices yielded qualitatively similar results.

### Sleep recordings

All sleep recordings were performed in a natural holding room adjoining the larger natural arena. Each monkey was kept in this holding room for a few days prior to the recording day so that it could get acclimatized to the room. This holding room contained a tree log platform and window, was separated from the natural arena by a partition with a metal grill, through which the monkey could interact with his social partner. On the recording day, we fixed the wireless logger around 3:00 PM, and brought it to the room with the touchscreen workstation where it performed a fixation task involving food box images. After the fixation task, the monkey was transferred back into the holding room where it remained for the rest of the night. We removed the tree log platform before moving the monkey to the holding room to better visibility from the CCTV cameras. We turned off the lights around 6:30 PM and waited in an adjoining room for the monkey to sleep, while monitoring and recording its video via infrared-capable video cameras.

Once the monkey started to sleep (as evidenced by its Head Down state, as defined below), we started the video synchronization task in the behavior room as detailed above (not visible to the monkey), remotely started the wireless brain recording and recorded for ∼2 hours in each monkey (121 min for M1, 135 minutes for M2), until the capacity limit of the wireless logger was reached, after which we stopped the recording remotely. Following this, we turned the lights back on around 10:00 PM and got the monkey back into the chair, snout-restrained him, retrieved the wireless logger, replaced the logger with the nightcap, and returned him back to the holding room.

#### Manual annotation of sleep videos

We manually annotated the CCTV video recordings of each monkey’s sleep session. Bouts of sleep and wakefulness were marked based on visible behavioral markers. During sleep, the monkey took up a huddled position while sitting on the floor with his head bent down. We annotated the start of assuming this position as “Head Down”. When he awoke, he opened his eyes, raised his head, moved around the holding room and made various body movements. We annotated the start of this behavior as “Head Up”. During data analyses, we binned the data into 1 s bins. Bins that contained a transition between these two states were excluded from analyses.

#### Artefact removal

We observed high-amplitude artefacts in the wideband data. These were high amplitude square waves, which occurred across all channels roughly in 44-minute intervals, and lasted for exactly 1.31 s. This characteristic made manual demarcation of these artefacts straightforward. The data was cleaned manually by labelling these artefacts and setting these values to NaN in the analyses.

### Decoding of sleep and wake states

To assess whether neural activity at any given time contains information about the sleep or wake states, we performed a linear decoding analysis. We took the firing rates across all channels in each 1 s time bin, and obtained population vectors for all time bins with an unambiguous sleep/wake label (i.e. excluding transitions). These population vectors along with their labels were used to train a linear decoder (*classify* function, MATLAB 2024). To evaluate the decoder, we performed a 5-fold cross-validation: we shuffled all the bins and divided them into 5 equal parts. For each held out 20% of the bins, we trained the decoder on the remaining 80% of the bins and tested on these held-out bins. In this manner we obtained a predicted label for every time bin.

### Exclusion of REM epochs close to the beginning of a sleep bout

Sleep epochs, which are at the beginning of a sleep bout, are likely a transition from wakefulness to sleep, and thus the neural activity pattern can be similar to the wake state. In addition, REM epochs are known to occur after an NREM phase/phase of deep sleep. Hence, we chose to exclude any candidate pREM points that occur too soon into the beginning of a sleep bout (a minimum duration of 200 s into the sleep bout was required to consider a bout as pREM). On the other hand, awakenings commonly occur right after a REM phase (Klemm, 2011). Hence, we did not remove the pREM points identified at the end of a sleep bout.

## ACKNOWLEDGEMENTS

We thank Dr. Ashesh Dhawale for valuable discussions, Dr. Tim Blanche of White-Matter LLC for technical support with wireless recordings, Dr. Sebastian Chandu for assistance with implant surgeries, Dr. Sumedh Shastry for help with physiology during surgery and post-surgical care, Dr. Sumedh Shastry & Mr. V. Ramesh for primate facility management and animal care and Mr. Ravi & Mr. Ashok for animal care.

## AUTHOR CONTRIBUTIONS

Jason Joby: Experiment Design, Data Collection, Curation & Analysis, Manuscript writing. Specific Contributions: designed experiment, collected data, performed data analysis, wrote manuscript sections.

Sini Simon: Recording setup, Analysis, Manuscript writing & editing. Specific contributions: Developed hardware & software for wireless brain recordings, tested electrodes, trained monkeys, planned and conducted implant surgeries, participated in troubleshooting recordings, reviewed data analysis, wrote manuscript sections and integrated comments from all authors.

Surbhi Munda: Recording setup, Experiment Design, Data collection. Specific contributions: Developed hardware & software for wireless brain recordings, trained monkeys, planned and conducted implant surgeries, participated in troubleshooting recordings, designed experiments, collected data, helped with data visualization, provided feedback on the manuscript drafts.

Shubhankar Saha: Recording setup. Specific contributions: Developed hardware & software for wireless brain recordings, trained monkeys, planned and conducted implant surgeries, provided feedback on manuscript drafts.

Thomas Cherian: Recording setup. Specific contributions: Developed hardware & template software codes for recording, tested electrodes and implant housing, planned and conducted implant surgeries, participated in troubleshooting recordings.

Georgin Jacob: Recording setup. Specific contributions: Developed hardware & template software codes for recording, developed software for automated spike detection and sorting, tested electrodes and implant housing, planned and conducted implant surgeries, participated in troubleshooting recordings, and provided feedback on manuscript drafts.

Jhilik Das: Recording setup, Manuscript editing. Specific contributions: Developed hardware & software codes for data curation and analysis, tested electrodes and implant housing, snout trained monkeys, planned and conducted implant surgeries, participated in troubleshooting recording setup, and provided feedback on manuscript drafts.

Debdyuti Bhadra: Recording setup, Data collection. Specific contributions: Designed experiments, collected data, helped with hardware and software updates for wireless brain recordings, planned and conducted implant surgeries, provided feedback on manuscript drafts.

SP Arun: Funding, Project Management, Experiment Design, Recording Setup, Data Collection, Curation & Analysis, Manuscript writing. Specific Contributions: supervised hardware & software development, electrode testing & implant testing, performed implant surgeries, participated in troubleshooting recordings, designed experiment, collected data, supervised data analysis, wrote manuscript.

## DATA AVAILABILITY

All data and code required to reproduce the results are publicly available at https://osf.io/2gfrd

## SUPPLEMENTARY MATERIAL

**Figure S1.**
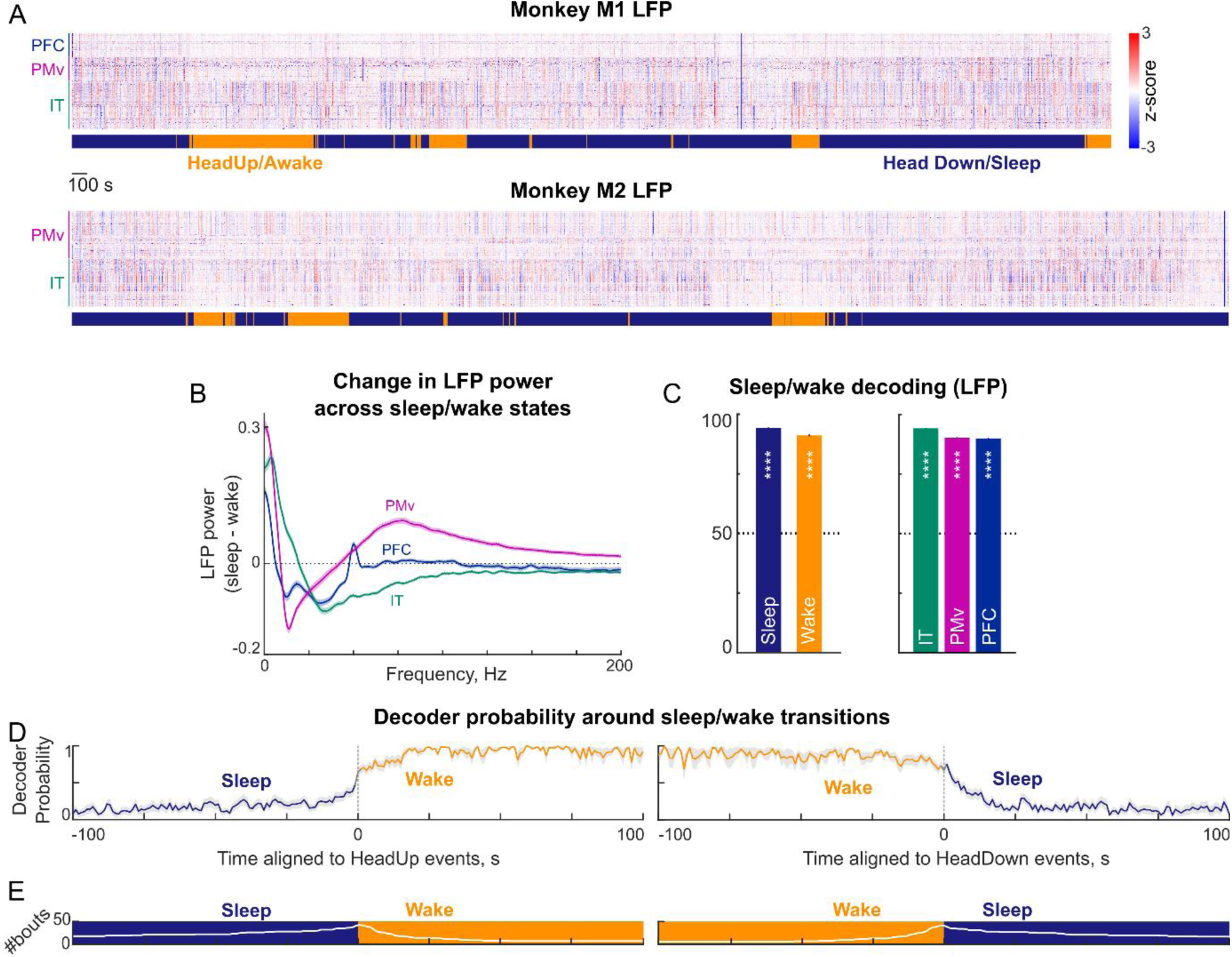
Neural signatures of sleep in the local field potential (LFP) (A) LFP amplitude (z-scored, in 1 ms bins) across all recorded sites with periods of wake and sleep indicated below for monkey M1 (*top*) and M2 (*bottom*). (B) Difference in LFP spectral power between sleep and wake states plotted against frequency (mean ± s.e.m across neurons) for IT (*green*), PMv (*purple*) and PFC (*blue*) recording sites. LFP spectral power was calculated in 1 s bins (pspectrum function, MATLAB 2024), represented in log10 units and then averaged across all bins of a given state. Positive values indicate higher power during sleep. IT showed increased power in 0-19Hz, and decreased power in >19Hz. PFC showed increased power in 0-6Hz and decreased power in >6Hz. By contrast, PMv showed increased power in both low (0-9Hz) and high frequencies (>43 Hz) during sleep. (C) Decoding of sleep/wake state. *Left:* Decoding accuracy (mean ± s.e.m across bins) of a linear decoder trained on LFP spectral power (0-200 Hz) from all neurons on sleep/wake states, pooled across monkeys. *Right:* Decoding accuracy separately for each brain region. (D) Decoder probability (mean ± s.e.m across bouts) for sleep/wake states averaged around the time of transitions from sleep to wake (*left*) and wake to sleep (*right*). (E) Number of bouts contributing at each time to the averaging in panels (D) and (E), for sleep to wake (*left*) and wake to sleep (*right*) transitions.

**Figure S2.**
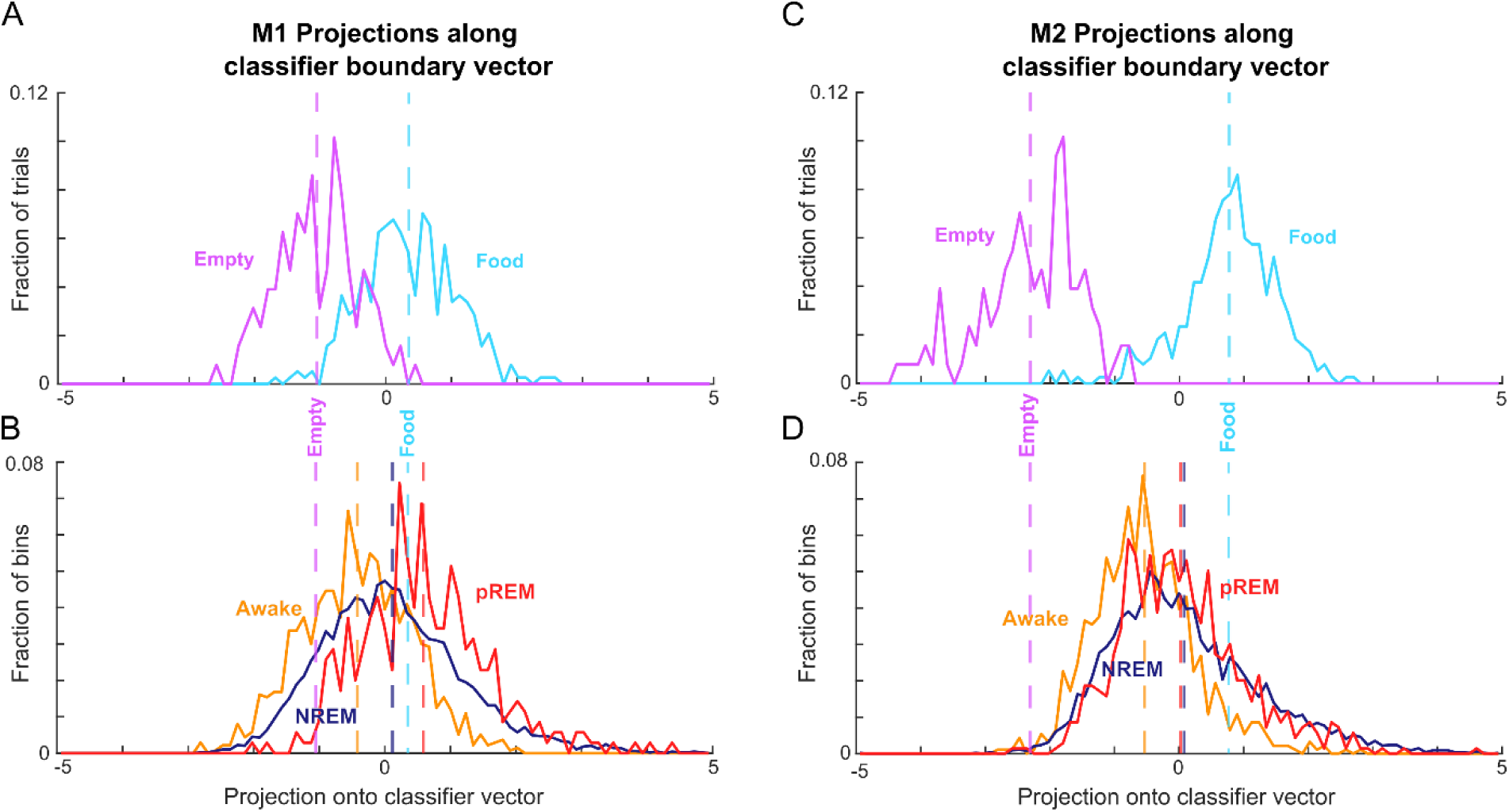
Food classifier projections while viewing images and during sleep. (A) Histogram of classifier projections of the food decoder trained on M1 IT neural responses in the fixation task, for trials with images of the food box without food (*purple*) and with food (*cyan*). (B) Histogram of classifier projections of the food decoder trained on the fixation task but tested on the sleep recording of monkey M1 for NREM bins (n = 4875), pREM bins (n = 350) and Awake bins (n = 1367). It can be seen that classifier projections are in the same range as the projections in the fixation task, confirming that neural responses during sleep are not qualitatively different. (C) Same as (A), but for monkey M2 NREM bins (n = 5443), REM bins (n = 703) and Awake bins (n = 932). (D) Same as (B), but for monkey M2. It can be seen that classifier projections are in the same range as the projections in the fixation task, confirming that neural responses during sleep are not qualitatively different.

**Figure S3.**
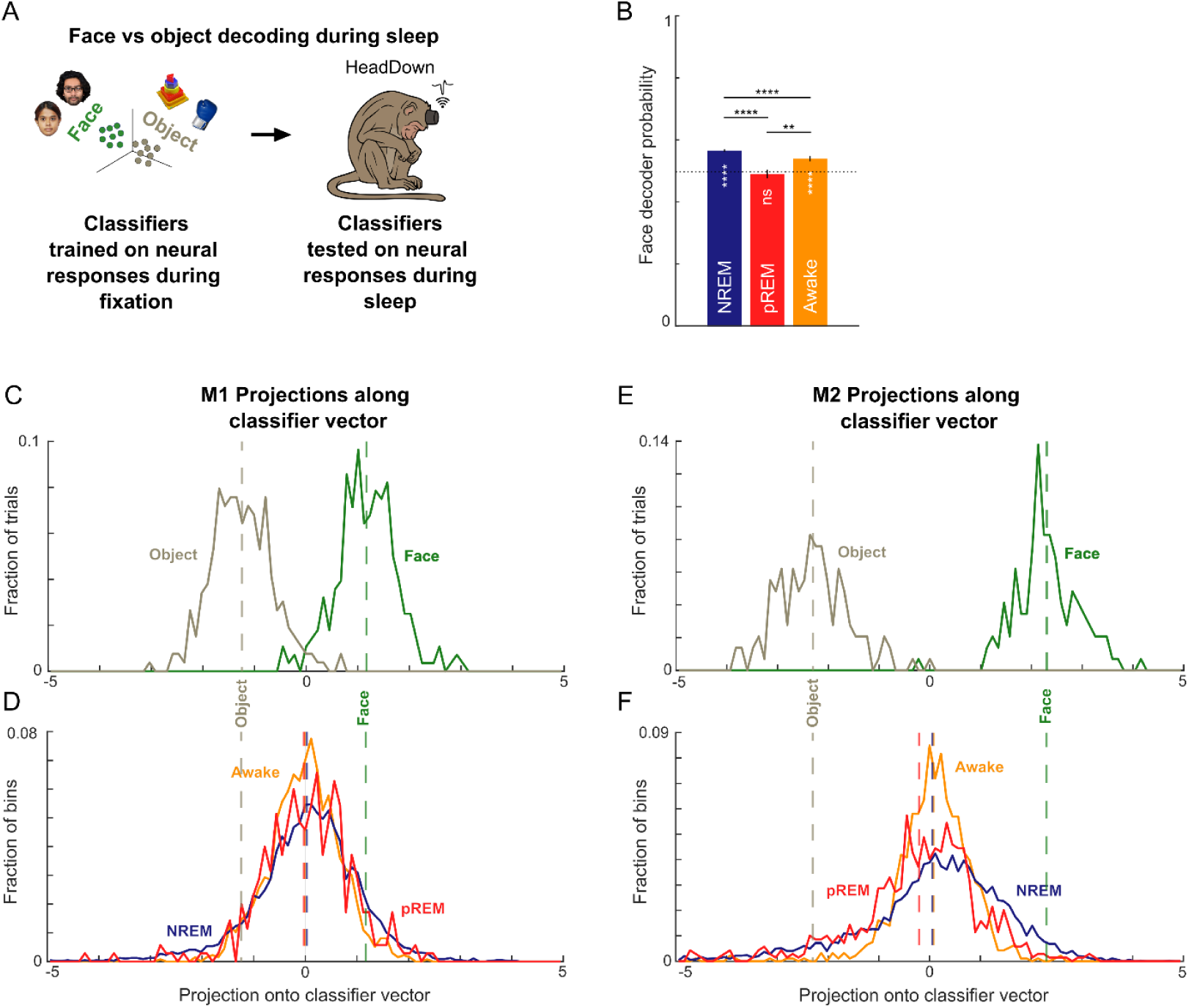
Face classifier projections while viewing images and during sleep. (A) Schematic of face decoding: a linear decoder was trained on neural responses in IT to objects and faces, and then tested on neural activity during sleep. *[Face images shown are those of the authors J.D. and S.S.]* (B) Decoder probability (mean ± s.e.m) across the NREM, pREM and Awake bins pooled across monkeys. Asterisks spanning bars indicate statistical significance of the comparison between states (**** is p < 0.0005, rank-sum test comparing decoder probability across time bins). Asterisks within bars indicate statistical significance compared to chance (50%) (**** is p < 0.0005, sign-rank test comparing decoder probability across time bins with that expected by chance) (C) Histogram of classifier projections of the food decoder trained on M1 IT neural responses in the fixation task, for trials with images of the objects (*light brown*) and faces (*green*). (D) Histogram of classifier projections of the food decoder trained on the fixation task but tested on the sleep recording of monkey M1 for NREM bins (n = 4875), pREM bins (n = 350) and Awake bins (n = 1367). It can be seen that classifier projections are in the same range as the projections in the fixation task, confirming that neural responses during sleep are not qualitatively different. (E) Same as (A), but for monkey M2 NREM bins (n = 5443), pREM bins (n = 703) and Awake bins (n = 932). (F) Same as (B), but for monkey M2. Note that classifier projections are in the same range as the projections in the fixation task, confirming that neural responses during sleep are not qualitatively different.

